# Predicting cellular position in the *Drosophila* embryo from Single-Cell Transcriptomics data

**DOI:** 10.1101/796029

**Authors:** Jovan Tanevski, Thin Nguyen, Buu Truong, Nikos Karaiskos, Mehmet Eren Ahsen, Xinyu Zhang, Chang Shu, Ke Xu, Xiaoyu Liang, Ying Hu, Hoang V.V. Pham, Li Xiaomei, Thuc D. Le, Adi L. Tarca, Gaurav Bhatti, Roberto Romero, Nestoras Karathanasis, Phillipe Loher, Yang Chen, Zhengqing Ouyang, Disheng Mao, Yuping Zhang, Maryam Zand, Jianhua Ruan, Christoph Hafemeister, Peng Qiu, Duc Tran, Tin Nguyen, Attila Gabor, Thomas Yu, Enrico Glaab, Roland Krause, Peter Banda, DREAM SCTC Consortium, Gustavo Stolovitzky, Nikolaus Rajewsky, Julio Saez-Rodriguez, Pablo Meyer

**Author notes:** These authors contributed equally (Jovan Tanevski, Thin Nguyen, Buu Truong). DREAM SCTC Consortium authors and affiliations are listed in the supplementary material.

## Abstract

Single-cell RNA-seq technologies are rapidly evolving but while very informative, in standard scRNAseq experiments the spatial organization of the cells in the tissue of origin is lost. Conversely, spatial RNA-seq technologies designed to keep the localization of the cells have limited throughput and gene coverage. Mapping scRNAseq to genes with spatial information increases coverage while providing spatial location. However, methods to perform such mapping have not yet been benchmarked. To bridge the gap, we organized the DREAM Single-Cell Transcriptomics challenge focused on the spatial reconstruction of cells from the *Drosophila* embryo from scRNAseq data, leveraging as gold standard genes with *in situ* hybridization data from the Berkeley *Drosophila* Transcription Network Project reference atlas. The 34 participating teams used diverse algorithms for gene selection and location prediction, while being able to correctly localize rare subpopulations of cells. Selection of predictor genes was essential for this task and such genes showed a relatively high expression entropy, high spatial clustering and the presence of prominent developmental genes such as gap and pair-ruled genes and tissue defining markers.

## 1 Introduction

The recent technological advances in single-cell sequencing technologies have revolutionized the biological sciences. In particular single-cell RNA sequencing (scRNAseq) methods allow transcriptome profiling in a highly parallel manner, resulting in the quantification of thousands of genes across thousands of cells of the same tissue. However, with a few exceptions [1, 2, 3, 4, 5] current high-throughput scRNAseq methods share the drawback of losing the information relative to the spatial arrangement of the cells in the tissue during the cell dissociation step.

One way of regaining spatial information computationally is to appropriately combine the single-cell RNA dataset at hand with a reference database, or atlas, containing spatial expression patterns for several genes across the tissue. This approach was pursued in a few studies [6, 7, 8, 9, 10]. Achim *et al* identified the location of 139 cells using 72 reference genes with spatial information from whole mount *in situ* hybridization (WMISH) of a marine annelid and Satija *et al* developed the *Seurat* algorithm to predict position of 851 zebrafish cells based on their scRNAseq data and spatial information from *in situ*-hybridizations of 47 genes in ZFIN collection [11]. In both cases, cell positional predictions stabilized after the inclusion of 30 reference genes. Karaiskos *et al* reconstructed the early *Drosophila* embryo at single-cell resolution and while the authors were successful in their reconstruction, their approach did not lead to a predictive algorithm and mainly centered around maximizing the correlation between scRNAseq data and the expression patterns from *in situ*-hybridizations of 84 mapped genes in The Berkeley *Drosophila* Transcription Network Project (BDTNP). In this project, *in situ* hybridization data was collected resulting in a quantitative high-resolution gene expression reference atlas [12]. Indeed, Karaiskos *et al* showed that the combinatorial expression of these 84 BDTNP markers suffice to uniquely classify almost every cell to a position within the embryo.

In the absence of a reference database, it is also possible to regain spatial information computationally solely from the transcriptomics data by leveraging general knowledge about statistical properties of spatially mapped genes against the statistical properties of the single-cell RNA dataset [13, 14]. Bageritz *et al.* were able to reconstruct the expression map of a *Drosophila* wing disc using scRNAseq data by correlation analysis. They exploited the coexpression of non-mapping genes to a few mapping genes with known expression patterns, to predict the spatial expression patterns of 824 genes [13]. Nitzan *et al.* exploited the knowledge of the distribution of distances between mapping genes in physical space to predict the possible locations of cells based on the distribution of distances between genes in the expression space. Following this approach, they were able to successfully reconstruct the locations of cells of the *Drosophila* and zebrafish embryos from scRNAseq data [14]. Although these approaches have indicated important steps to reconstruct the position of a cell in a tissue from their RNAseq expression, a global assessment is needed to evaluate the methods used and the number and nature of the genes with spatial expression information required for correctly assigning a location to each cell.

With this purpose in mind, and to catalyze the development of new methods to predict the location of cells from scRNAseq data we organized the DREAM Single cell transcriptomics challenge which ran from September through November 2018. DREAM challenges are a platform for crowdsourcing collaborative competitions[15] where a rigorous evaluation of each submitted solution allows for the comparison of their performance. The quality and reproducibility of each provided solution is also ensured. The combination of the individual solutions, i.e., the different approaches and insights to a common problem, leads to an overall wisdom-of-the-crowds (WOC) solution, with generally superior performance to any individual solution, from where collective insights can be garnered. We set up the challenge with 3 goals in mind. First, we used the data from Karaiskos *et al* to foster the design of a variety of algorithms and objectively tested how well they could predict the localization of the cells. Second, we evaluated how the predictive performance of the algorithms was impacted by the number of reference genes from BDTNP with *in situ* hybridization information included in the predictions. Third, we investigated how the biological information carried in the selected genes was implemented in the algorithms to determine embryonic patterning.

The challenge, a first of its kind for single cell data, consisted of predicting the position of 1297 cells among 3039 *Drosophila melanogaster* embryonic locations for one half of a stage 6 pre-gastrulation embryo from their scRNAseq data (Figure 1A) [10]. At this stage cells in the embryo are positioned in a single two dimensional sheet following a bilateral symmetry, so that only positions in one half of the embryo where considered - accounting for the 3039 locations. Participants used the scRNAseq data for each of the 1297 cells obtained from the dissociation of 100-200 stage 6 embryos and the spatial expression patterns from *in situ*-hybridizations of 84 genes in the BDTNP database [12]. Gene determinants of different tissues such as neurectoderm, dorsal ectoderm, mesoderm, yolk and pole cells were provided as a hint. To aid the development of prediction algorithms, we provided (when available) the regulatory relationship -positive or negative-between the 84 genes in the *in situ*-hybridizations and the rest of the genes. We asked participants to provide an ordered list of 10 most probable locations in the embryo predicted for each of the 1297 cells using the expression patterns from (i) 60 genes out of the 84 in subchallenge 1, (ii) 40 genes out of the 84 in subchallenge 2, and (iii) 20 genes out of the 84 in subchallenge 3. The predictions were compared to the ground truth location determined by calculating the maximum correlation using all 84 *in situs* [10]. We received submissions from 34 teams, and the overall analysis of the results showed that the selection of genes is essential for accurately locating the cells in the embryo. The most selected genes had a relatively high expression entropy, showed high spatial clustering and featured developmental genes such as gap and pair-ruled genes in addition to tissue defining markers.

**Figure 1:**
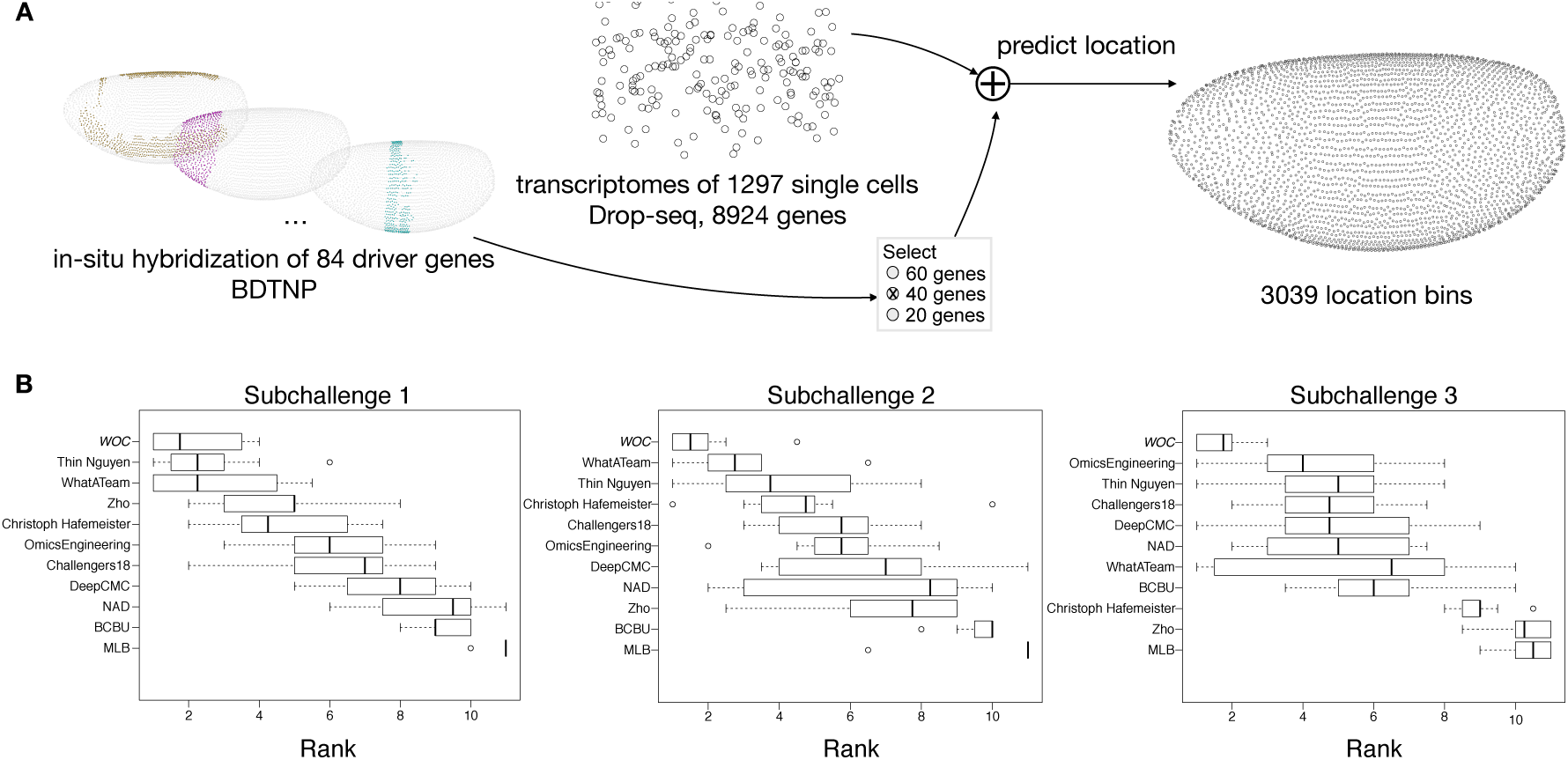
Overview of the challenge and results. **A.** In the DREAM Single-Cell Transcriptomics Challenge participants were asked to map the location of 1297 cells to 3039 location bins of an embryo of *Drosophila melanogaster*, by combining the scRNAseq measurements of 8924 genes for each cell and the spatial expression patterns from *in situ* hybridization of 60, 40 or 20 genes, for subchallenge 1, 2 and 3 respectivelly, for each embryonic location bin, selected from a total of 84 driver-genes. **B.** Ranking of the top 10 best performing teams and a wisdom of the crowds (WOC, in italic) solution, based on results from a post challenge cross-validated selection and prediction performance measured with three complementary scoring metrics. The boxplots show the distribution of ranks for each team on the 10 test folds. The rank for each fold is calculated as the average of the ranking on each scoring metric.

## 2 Results

### 2.1 Challenge setup

A distinctive feature of the single cell transcriptomics challenge was the public availability of the entire dataset and the ground truth locations produced by DistMap, a method using the *in situ*-hybridizations available at BDTNP [12], published together with the data [10]. We took three actions to mitigate the issue of not having a blinded ground truth. First, for the purpose of predictor gene selection, we allowed the use of scRNA-Seq data and biological information from other databases, but prohibited the use of *in situ* data. Second, to assess the quality of predictions, we devised three scores (detailed in the Methods section) that were not disclosed to the participants during the challenge. The scores measured not only the accuracy of the predicted location, but also how well the expression in the cell at the predicted location correlates with the expression from the reference atlas, the variance of the predicted locations for each cell, and how well the gene-wise spatial patterns were reconstructed. Finally, we devised a post-challenge cross-validation scheme to evaluate the soundness and robustness of the methods.

The challenge was organized in two rounds, a leaderboard round, and a final round. During the leaderboard round the participants were able to obtain scores for five submitted solutions before submitting a single solution in the final round. We received submissions from 40 teams in the leaderboard round and 34 submissions in the final round. Out of the 34 teams that made submissions in the final round, 29 followed up with public write-ups of their approaches and source code. For subchallenges 1 and 3 we were able to determine a clear best performer, but for subchallenge 2, there were two top ranked teams with statistically indistinguishable difference in performance (see Supplementary Figures S1,S2 and S3).

As stated, given that the ground truth for this challenge was publicly available and to avoid over-fitting, we decided to invite the top 10 performing teams to contribute to a post-challenge collaborative analysis phase to assess the soundness and stability of their gene selection and cell location prediction. Consequently, teams were tasked to provide predictions for a 10-fold cross-validation (CV) scenario, under the same conditions as for the challenge phase. The folds were extracted from the same RNA-seq dataset as in the challenge and every team used the same assignment of cells to folds. We evaluated the performance of the teams using the same scoring approach as in the challenge. To ensure the validity of the findings we decided to perform all further analysis and interpretation only from the results of the post-challenge phase.

### 2.2 Overview of results

Interestingly, for subchallenge 1 and 2, when participants had to use 60 or 40 genes for their predictions, the ranking of the best performing teams in the CV scenario did not change significantly compared to the challenge (Figure 1B cf. Figures S1 and S2). This was not the case in subchallenge 3 as no particular team from the top 10 outperformed in a statistically significant way the others when using 20 genes for their predictions. The results from the cross-validation showed that the approaches generalize well, i.e. the gene selection is performed consistently across the folds and the variance of the achieved scores across the folds is small for all teams (Figure S4). For each subchallenge we combined the gene selection and location predictions from the top 10 participants into a WOC solution (see details below) that performed better compared to the individual solutions (Figure 1B). The scores obtained by the best performing teams and the WOC solution are shown in Table 1.

**Table 1:** Best mean score for metrics *s*_1_, *s*_2_ and *s*_3_ achieved by the teams (Thin Nguyen, WhatATeam and OmicsEngineering) and the WOC solution. The standard deviation of scores across folds are in parenthesis. For more details on the scoring metrics see the Methods section.

| | $s_1$ | | $s_2$ | | $s_3$ | |
| --- | --- | --- | --- | --- | --- | --- |
|  | Teams | WOC | Teams | WOC | Teams | WOC |
| Subchallenge 1 | **0.76**($\pm 0.04$) | 0.73($\pm 0.04$) | **2.52**($\pm 0.28$) | 2.16($\pm 0.20$) | 0.59($\pm 0.01$) | **0.62**($\pm 0.01$) |
| Subchallenge 2 | 0.69($\pm 0.03$) | **0.70**($\pm 0.05$) | 1.16($\pm 0.12$) | **1.84**($\pm 0.26$) | **0.67**($\pm 0.02$) | 0.65($\pm 0.01$) |
| Subchallenge 3 | 0.65($\pm 0.05$) | **0.68**($\pm 0.03$) | 0.88($\pm 0.13$) | **1.42**($\pm 0.16$) | **0.79**($\pm 0.02$) | 0.71($\pm 0.01$) |

A summary of the methods used by participants for gene selection and location prediction can be seen in Table S2. The most frequently used method by participants for location prediction was a similarity based prediction, such as the maximum Matthews correlation coefficient between the binarized transcriptomics and the *in situs* that was proposed by Karaiskos et al. [10]. Another well performing approach was combining the predictions of a machine learning model and the Matthews correlation coefficients. The models were trained to predict either the coordinates of each cell or the binarized values of the selected *in situs* given transcriptomics data as input. The predictions were then made by selecting the location bins that corresponded to the nearest neighbors of the predicted values.

The most frequently used method by participants for gene selection was unsupervised or supervised feature importance estimation and ranking. For example, in a supervised feature importance estimation approach a machine learning model is trained to predict the coordinates of each cell, given the transcriptomics data at input, that is, the genes with available *in situ* hybridization measurements or all genes. Different machine learning models were trained such as Random Forest (BCBU, OmicsEngineering) or a neural network (DeepCMC, NAD). There were examples of unsupervised feature importance estimation and ranking by expression based clustering (NAD, Christoph Hafemeister, MLB), or a greedy feature selection based on predictability of expression from other genes (WhatATeam). Background knowledge about location specific marker genes, or the expected number of location clusters, was used by a small number of teams (WhatATeam and NAD) to inform the gene selection. Given the diversity of approaches to gene selection, we focused our analysis on better understanding the properties of frequently selected genes and providing recommendations for future experimental designs.

### 2.3 Analysis of the location prediction

A recurrent observation across DREAM challenges is that an ensemble of individual predictions performs usually better and is more robust than any individual method [16, 17]. This phenomenon, common also in other contexts, is denoted as the wisdom-of-the-crowds (WOC) [15]. In a typical challenge, individual methods output a single probability reflecting the likelihood of occurrence of an event. The WOC prediction is then constructed in an unsupervised manner by averaging the predictions of individual methods.

Given that in the single cell RNAseq prediction challenge participants had to submit 10 positions per cell, we developed a novel method that is based on k-means clustering to generate the WOC predictions. A diagram of the k-means approach is given in Figure 2 where for each single cell we first used k-means clustering to cluster the locations predicted by the individual teams [18] where the euclidean distance between the locations was used as the distance metric. In order to find the optimal *k*, we used the elbow method, i.e. we chose a *k* that saturates the sum of squares between clusters [19]. Note that each cluster consists of a group of locations and each location is predicted by one or more teams. Hence, for each cluster we calculated the average frequency that its constituent locations are predicted by individual teams. We then picked the cluster with the highest average frequency as our final cluster and ranked each location in this cluster based on how frequently it was predicted by individual methods. For each cell, the final prediction of the proposed WOC method consisted of the top 10 locations based on the above ranking. The k-means approach is based on the intuition that a single cell belongs to one location and its expression is mostly similar to that of cells in locations surrounding it.

**Figure 2:**
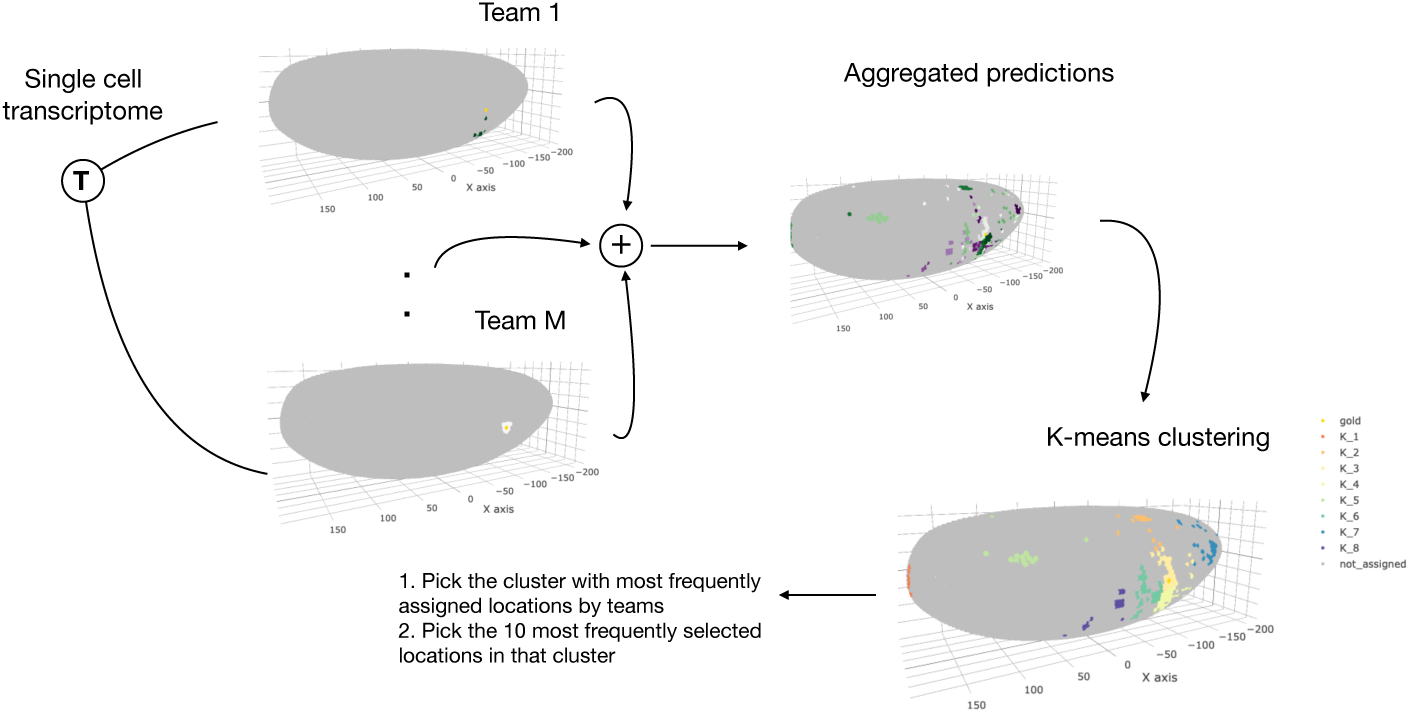
Wisdom of crowds location prediction. The location predictions for each cell by the top performing teams in the post-challenge cross-validation phase were aggregated in a wisdom of the crowds solution based on a k-means clustering approach.

The WOC location prediction approach does not take the genes used by the teams to make the predictions into account. However, after the WOC predictions are generated, in order to score them, we needed a list of genes for every subchallenge. To this end we used a WOC approach to gene selection (see the following section for more details) and used the most frequently selected genes per challenge. As reported above, the WOC solution performed better compared to the individual solutions (Figure 1B).

### 2.4 Analysis of selected genes

The selection of a subset of *in situs* used for cell location prediction was the hallmark that differentiated the subchallenges. It is unfeasible to evaluate all subsets of 20, 40 or 60 genes from the 84 due to the immense number of possible combinations of genes. Different approaches and heuristics can be used to select a subset of genes and the most frequent among the top 10 ranked teams were based on model based feature ranking algorithms, using normalized transcriptomics data (for more details see Table S2). However, if a subset of genes is selected as a candidate for solving the general task of location prediction, it should be consistently identified when similar sets of single cells are used as inputs. Therefore, we analyzed the consistency of gene selection for each team across folds by 10-fold cross-validation. More importantly, we were interested in subsets of genes that were consistently selected by multiple teams as this could underlie biological relevance.

The approaches for selecting genes taken by the top 10 teams resulted in consistent selection across folds, significantly better than random, for all subchallenges. Indeed, all of the pairwise Jaccard similarities of sets of selected genes for all teams were significantly higher than the expected Jaccard similarity of a random pair of subset of genes (see Supplementary Figure S4). Importantly, we measured an observable increase in variance and decrease of mean similarity as the number of selected genes decreased.

For each subchallenge we counted the number of times that the genes were selected by all teams in all folds. The genes, ordered by the frequency of selection in all subchallenges are depicted in Figure 3A. Forty percent of the top 20, 67% of the top 40 and 81% of the top 60 most frequently selected genes are the same for all three subchallenges (Figure 3B). The ranks assigned to all genes in the three subchallenges are highly correlated. Namely, the rank correlations range from 0.69 between subchallenges 1 and 3, to 0.83 between subchallenges 1 and 2, and subchallenges 2 and 3. Figure 3C shows a plot of the Jaccard similarity of the sets of top-k most frequently selected genes for pairs of subchallenges. We observe that a high proportion of genes are consistently selected across subchallenges. The lists of most frequently selected 60, 40 and 20 genes in subchallenges 1, 2 and 3 respectively are available in the supplementary material (Table S3).

**Figure 3:**
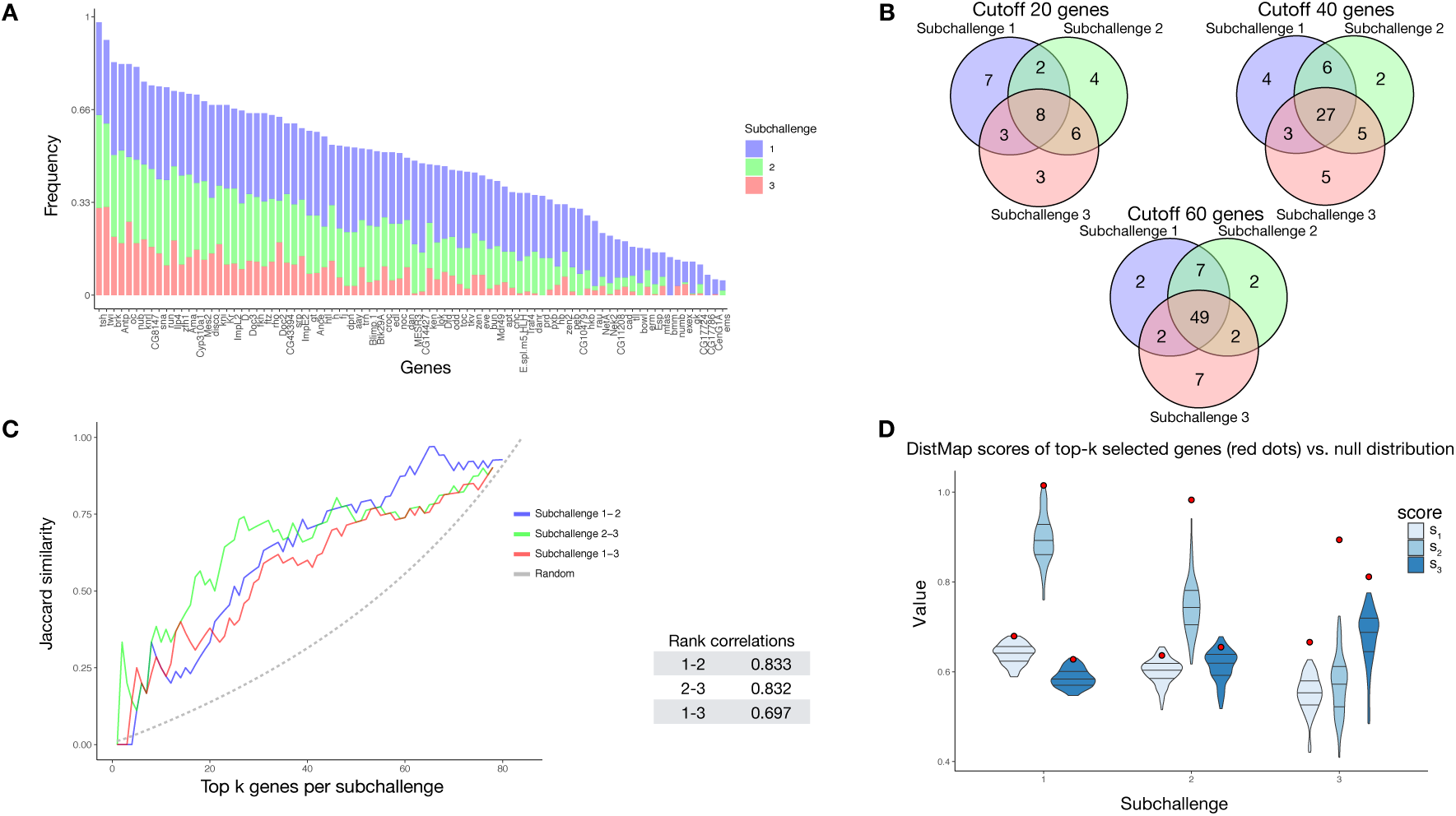
Analysis of gene selection. The results in all figures were generated from the genes that were selected by the top performing teams in the post-challenge cross-validation scenario. **A.** Frequency of selected genes in subchallenge 1 (blue), subchallenge 2 (green) and subchallenge 3 (red). The genes are ordered according to their cumulative frequency. **B.** Venn diagrams of the most frequently selected genes in the subchallenges with cutoff at 20, 40 and 60 most frequently selected genes, corresponding to the number of genes required for each subchallenge **C.** *Left*, the similarity of most frequently selected genes for pairs of subchallenges. The Jaccard similarity measures the ratio of the size of the intersection and the union of two sets 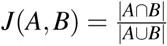 *Right*, table of correlations between gene rankings (by frequency) for pairs of subchallenges. **D.** Validation of the performances of the wisdom of the crowds (WOC) selection of genes, i.e the most frequently selected 60, 40 and 20 genes in the respective subchallenges. The violin plots represent null **distribution of scores obtained by 100 randomly selected sets of 60, 40 and 20 genes using DistMap**. The red dots represents the performance obtained by using DistMap with the WOC selection of genes.

We conclude that the gene selection is not only consistent by team across folds, but also across teams and subchallenges. This finding outlines a direction for further analysis, namely the validation of the predictive performance and analysis of the common properties of the most frequently selected genes.

#### 2.4.1 Validation of frequently selected genes

We defined a simple procedure to obtain a WOC gene selection for each of the subchallenges. It consisted on selecting the most frequently selected genes for each subchallenge (different colored bars in Figure 3A). For example, for subchallenge 1 we chose the 60 most frequently selected genes looking only at the heights of blue portion of the bar. Interestingly, the 20 most frequently selected genes in subchallenge 3 are included in the list of 40 most selected genes in subchallenge 2 (except for *Doc2*), conversely included in the list of 60 most selected genes in subchallenge 1.

To validate the predictive performance of the WOC gene selection, we predicted the cell locations using DistMap and scored the predictions using the same scoring metrics as for the challenge, estimating the significance of the scores through generated null distributions of scores for each subchallenge. The null distribution of the scores was generated by scoring the DistMap location prediction using 100 different sets of randomly selected genes. For each subchallenge and each score we estimated the empirical distribution function and then calculated the percentile of the values of the scores obtained with the WOC gene selection.

The null distributions and the values of the scores obtained with the WOC gene selection are shown in Figure 3D. All values of the scores for subchallenge 1 fall in the 99th percentile. For subchallenge 2 *s*_1_ and *s*_3_ fall into the 92nd percentile and *s*_2_ in the 100th percentile. For subchallenge 3 all scores fall in the 100th percentile. Overall the performance of DistMap with the WOC selected genes performs significantly better than a random selection of genes. The actual values of the scores are on par with those achieved by the top 10 teams in the challenge.

#### 2.4.2 Properties of frequently selected genes

We conjectured that the most frequently selected genes should carry enough information content collectively to uniquely encode a cell’s location. Furthermore, genes should also contain location specific information, i.e. their expression should cluster well in space. To quantify these features, we calculated the entropy and the join count statistic for spatial autocorrelation of the *in situs* (see Figure 4A and Methods for description). We observed that most of the *in situ* genes have relatively high entropy as observed by the high density in the upper part of the plots and show high spatial clustering, i.e show values of the join count test statistic lower than zero.

**Figure 4:**
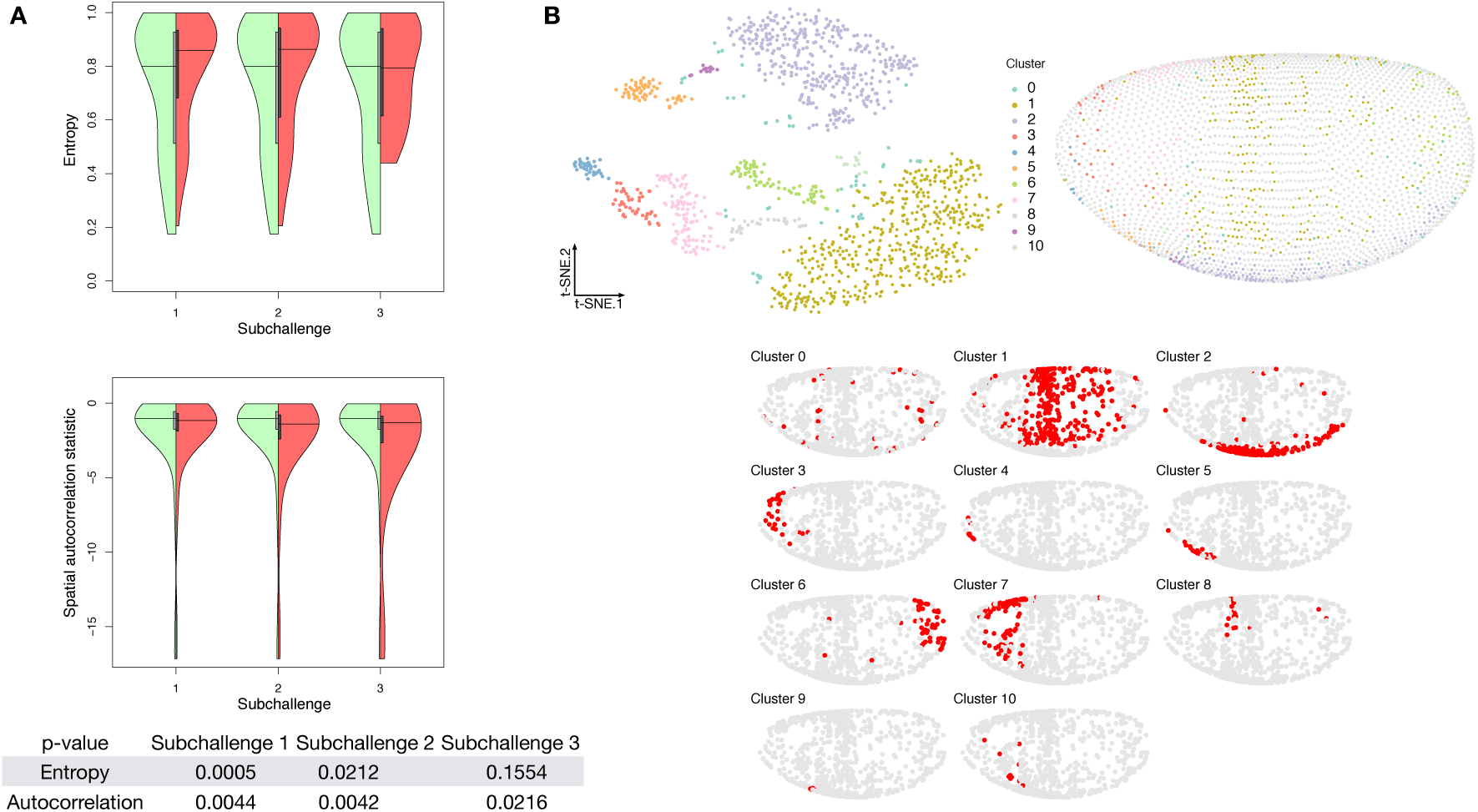
Properties of selected genes. **A.** Double violin plots of the distribution of entropy and spatial autocorrelation statistic of *Left, green* all *in situs* calculated on all embyonic location bins and *Right, red* the most frequently selected 60, 40 and 20 genes in the respective subchallenges. [bottom table] p-values of a one sided Mann-Whitney U test of location shift comparing the selected (red part of the violin plot) genes vs the non-selected genes. **B.** *Top left*, visualization of the transcriptomics data containing only the most frequently selected 60 genes from subchallenge 1 by the top performing teams (embedding to 2D by t-SNE). Each point (cell) is filled with the color of the cluster that it belongs to (density-based clustering with DBSCAN). *Top right*, spatial mapping of the cells in the *Drosophila* embryo as assigned by DistMap using only the 60 most frequently selected genes from subchallenge 1. The color of each point corresponds to the color of the cluster from the t-SNE visualization. *Bottom*, highlighted (red) location mapping of cells in the Drosophila embryo for each cluster separately.

To test our conjectures of high entropy and spatial correlation we tested the significance of the shift of the values between the WOC selected genes and the non-selected genes from all *in situs* for each subchallenge. Since the Shapiro-Wilk test of normality rejected the null-hypothesis for both entropy and join count metrics (*p <* 2.3 10^*-*6^ and *p <* 1.8 10^*-*15^) that their values are distributed normally for the *in situ* genes, we opted for a nonparametric, one sided Mann-Whitney U test. We observed significant value shift for the autocorrelation statistic for all subchallenges 1 to 3 (see bottom of Figure 4A right red part of violin plots and table). Although we see a decrease of the statistical significance of the mean value shift for the distribution of values of the entropy of the selected subsets of genes, the shift is significant for all subchallenges and at the same time, we observe that tail of the distribution shortens.

To test whether the information relative to different cell types is retained with the selected subset of 60, 40 or 20 WOC selected genes, we embedded the cells into 2D space using t-distributed stochastic embedding (t-SNE) [20] aiming for high accuracy (*θ* = 0.01), Figure 4B and Figure S5. We then clustered the t-SNE embedded data using density-based spatial clustering of applications with noise (DBSCAN) [21]. DBSCAN determines the number of clusters in the data automatically based on the density of points in space. The minimum number of cells in a local neighborhood was set to 10 and the parameter *ε* = 3.5 was selected by determining the elbow point in a plot of sorted distances of each cell to its 10th nearest neighbor. We found that the 9 prominent cell clusters identified in the study by Karaiskos *et al.* [10] are preserved in our t-SNE embedding and clustering experiments when considering the most frequently selected 60 or 40 genes from subchallenges 1 and 2. The number of clusters of cells with specific localization is reduced when considering the most frequently selected 20 genes from subchallenge 3.

We next associated the properties of the *in situs* that were found to be indicative of good performance in the task of location prediction with statistical properties of the genes in the transcriptomics data. Our goal was to discover statistical properties of the transcriptomics data that might inform future experimental designs when selecting target genes for *in situ* hybridizations. We calculated statistical features across cells for the subset of genes from the transcriptomics data for which we also have *in situ* measurements. These include the variance of gene expression σ^2^ across cells, the coefficient of variation 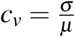, the number of cells with expression zero 0 and the entropy of binarized expression *H*_*b*_. We then calculated the correlation across genes for each of these metrics and the measured spatial properties of interest of the *in situs*, i.e entropy *H* and the value of the joint count statistic *Z* (see Table 2). Although the selection of highly variable genes was one of the approaches used by some of the top 10 teams, the variance for each gene in the scRNAseq expression, although highly correlated to the entropy of the corresponding *in situ* measurements of that gene, it is less correlated than other properties. Also, we observed that the positive correlation of the entropy to the variance of each gene, becomes a negative correlation against their coefficient of variation. This negative correlation can have two sources, the genes with high entropy may have low standard deviation or high mean expression. Since we observe positive correlation of entropy to the variance of expression, we can conclude that the negative correlation is a result of highly expressed genes. Since a known drawback of scRNAseq is a high number of dropout events for lowly expressed genes [22], this observation is further supported by the negative correlation of the entropy and the number of cells with zero expression. We observed the highest correlation of *in situ* entropy to the entropy of the binarized expression. Regarding the spatial autocorrelation, all statistical features of the transcriptomics were only slightly positively correlated to the join count statistic except for the entropy of binarized expression which had slightly negative correlation.

**Table 2:** Correlations of transcriptomics to *in situ* properties of the genes where both measurements are available. σ ^2^ - variance of a gene across cells, *c*_*v*_ - coefficient of variation, 0 - number of cells with zero expression, *H*_*b*_ - entropy of binarized expression, *H* - entropy, *Z* - join count test statistic

| | $\rho$ | *in situ* | |
| --- | --- | --- | --- |
| | | $H$ | $Z$ |
| scRNAseq | $\sigma^2$ | 0.50 | 0.18 |
| | $c_v$ | -0.69 | 0.26 |
|  | 0 | -0.64 | 0.29 |
| | $H_b$ | 0.72 | -0.30 |

## 3 Discussion

In this paper we report the results of a crowdsourcing effort organized as a DREAM challenge, around the issue of predicting the spatial arrangement of cells in a tissue from scRNAseq data. Analysis of the top performning methods and their performance provided us a number of unbiased insights. First, it unveiled a connection in the cell-to-cell variability in *Drosophila* embryo gene expression and the selection of the best genes for predicting the localization of a cell in the embryo from their scRNAseq expression. The most selected genes had a relatively high entropy, hence high variance and expression while also showing high spatial clustering. The smaller the number of selected genes, i.e going from subchallenge 1 to 3, the more these features became apparent. The observed advantage of genes with high overall expression in cells might lead to less dropout counts in the scRNAseq data, a known disadvantage of the technology, leading to more accuracy in the cell placement. We also found that the 9 prominent spatially distinct cell clusters previously identified are preserved when considering the most frequently selected 60 or 40 genes, but the number of clusters is reduced when considering only the most frequently selected 20 genes. This finding is in line with the conclusions of Howe et al. [11] where in a related task of location prediction the performance stabilized after the inclusion of 30 genes in a related experiment. The WOC gene selection and the k-means clustered WOC model for cell localization performed comparably or better than the participant’s models, showing once more the advantage of the wisdom-of-the-crowds. All these results can be explored in animated form at https://dream-sctc.uni.lu/.

Given that it has been shown that positional information of the anterior-posterior (A-P) axis is encoded as early in the embryonic development as when the expression of the gap genes occurs [23, 24], we thought that it should be possible to implement in algorithms for this challenge the information contained in the regulatory networks of *Drosophila* development [25]. Although only a small number of participants, among them the best performers, directly used biological information related to the regulation of the genes or their connectivity, the most frequently selected genes in all 3 subchallenges have interesting biological properties. Indeed, gap genes such as *giant* (*gt*), *kruppel* (*kr*), *knirps* (*kni*) were selected in all 3 subchallenges (see Figure 5 and Table S3 that also includes *kni*-like *knrl*) although *tailless* (*tll*) and *hunchback* (*hb*) were not. Along the A-P axis, maternally provided *bicoid* (*bcd*) and *caudal* (*cad*) first establish the expression patterns of gap and terminal class factors, such as *hb, gt, kr* and *kni*. These A-P early regulators then collectively direct transcription of A-P pair-rule factors, such as *even-skipped* (*eve*), *fushi-tarazu* (*ftz*), *hairy* (*h*), *odd skipped*, (*odd*), paired (*prd*) and runt (*run*) which in turn cross-regulate each other. Not being part of the *in situs*, neither *bcd*, nor *cad* were selected but *ama* sitting near *bcd* in the genome might have been selected for its similar expression properties. Furthermore, we also find that pair-rule genes were most prominently selected in subchallenges 1 (*eve, odd, prd*, the Paired-like *bcd* and *bcd*) and 2 (*h, ftz* and *run*). A similar cascade of maternal and zygotic factors controls patterning along the dorsal-ventral (D-V) axis were *dorsal* (*d*), *snail* (*sna*) and *twist* (*twi*) specify mesoderm and the pair rule factors *eve* and *ftz* specify location along the trunk of the A-P axis. Again, *sna* and *twi* were selected in all subchallenges and *d* in subchallenges 1 and 2. These selected transcription factors specify distinct developmental fates and can act via different cis-regulatory modules but their quantitative differences in relative levels of binding to shared targets correlates with their known biological and transcriptional regulatory specificities [26]. The rest of the selected genes were the homeobox genes (*nub, antp*) and differentiators of tissue such as mesoderm (*ama, mes2, zfh1*), ectoderm (*doc2* and *doc3*), neural tissue (*noc, oc, rho*) and EGFR pathway (*rho, edl*). The complete lists of most frequently selected genes are available in Table S3.

**Figure 5:**
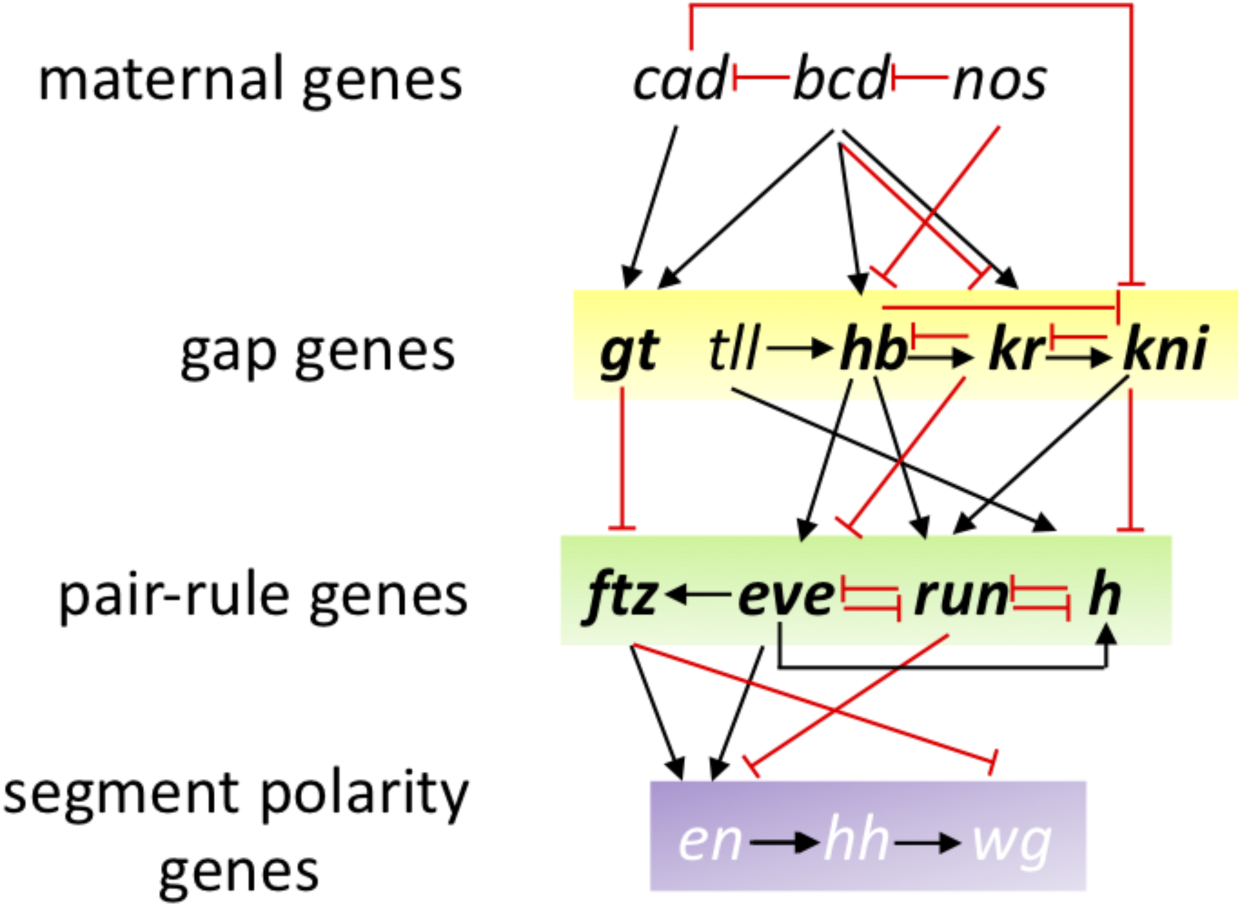
Gene regulatory network of early *Drosophila* development. Not all regulations are represented, nor pair ruled genes *odd* & *prd*. Frequently selected genes are represented in bold.

Since the ground truth of single cell locations was publicly available, the organization of this DREAM challenge brought risks that, given the importance of the scientific question asked, we thought worth taking. However, without the post-challenge phase it would have been impossible to distinguish the robust and sound methods from methods that were overfitting the results. Overall, the single cell transcriptomics challenges unveils not only the best gene-selection methods and prediction approaches to localize a cell in the *Drosophila* embryo, but also explains the biological and statistical properties of the genes selected for the predictions. Further identification of additional properties such as spatially autocorrelated genes might require the use of alternative scRNAseq focused approaches [13, 14]. However, we think that the approach defined here could be used or adapted when performing similar cell-placing tasks in other organisms, including human tissues. Given the importance of spatial arrangements for disease development and treatment, we foresee an application of these methods to medical questions as well.

## 4 Methods

### 4.1 Scoring

We scored the submissions for the three subchallenges using three metrics *s*_1_, *s*_2_ and *s*_3_.*s*_1_ measured how well the expression of the cell at the predicted location correlates to the expression from the reference atlas and included the variance of the predicted locations for each cell. While *s*_2_ measured the accuracy of the predicted location and *s*_3_ measured how well the gene-wise spatial patterns were reconstructed.

Let *c* represent the index of a cell, given in the transciptomics data in the challenge where 1 ≤ *c* ≤ 1297. Each cell *c* is located in a bin ε_*c*_∈ {1 ‥3039} at a position with coordinates *r*(ε_*c*_) = (*x*_*c*_, *y*_*c*_, *z*_*c*_). Each cell is associated with a binarized expression profile *t*_*c*_ = (*t*_*c*1_,*t*_*c*2_,…,*t*_*cE*_), where 1 ≤ E ≤ 8924, and a corresponding binarized *in situ* profile *f*_*c*_ = (*f*_*c*1_, *f*_*c*2_, …, *f*_*cK*_), where the maximum possible value of *K* for which we have *in situ* information is *K* = 84. For different subchallenges we consider *K* ∈ {20, 40, 60}. Using *K* selected genes the participants were asked to provide an ordered list of 10 most probable locations for each cell. We represent with the mapping function *A*(*c, i, K*) the value of the predicted *i*-th most probable location for cell *c* using *K in situs*. For the first scoring metric *s*_1_ we calculated the weighted average of the Mathews correlation coefficient (MCC) between the *in situ* profile of the ground truth cell location 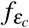 and the *in situ* profile of the most probable predicted location *f*_*A*(*c,*1,*K*)_ for that cell

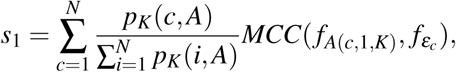

where *N* is the total number of cells with predicted locations.

The Matthews correlation coefficient, or *ϕ* coefficient, is calculated from the contingency table obtained by correlating two binary vectors. The MCC is weighted by the inverse of the distance of the predicted most probable locations to the ground truth location *p*_*K*_(*c*). The weights are calculated as 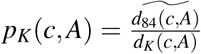, where 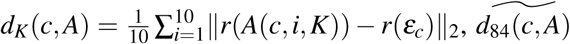 is the value of *d*_*K*_(*c, A*) using the ground truth most probable locations assigned with *K* = 84 using DistMap, and ‖ · ‖_2_ is the Euclidean norm.

The second metric *s*_2_ is simply the average inverse distance of the predicted most probable locations to the ground truth location

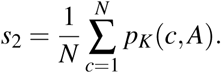

Finally, the third metric *s*_3_ measures the accuracy of reconstructed gene-wise spatial patterns

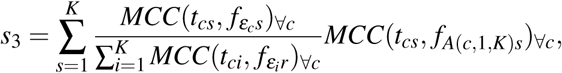

where ∀_*c*_ denotes that the *MCC* is calculated cell wise for each gene.

For 287 out of the 1297 cells, the ground truth location predictions were ambiguous, i.e., the MCC scores were identical for multiple locations. These cells were removed both from the ground truth and the submissions before calculating the scores.

The teams were ranked according to each score independently. The final assigned rank *r*_*t*_ for team *t* was calculated as the average rank across scores. Teams were ranked based on the performance as measured by the three scores on 1000 bootstrap replicates of the submitted solutions. The three scores were calculated for each bootstrap. The teams were then ranked according to each score. These ranks were then averaged to obtain a final rank for each team on that bootstrap. The winner for each subchallenge was the team that achieved the lowest ranks. We calculated the Bayes factor of the bootstrap ranks for the top performing teams. Bayesian factor of 3 or more was considered as a significantly better performance. The Bayes factor of the 1000 bootstrapped ranks of teams *T*_1_ and *T*_2_ was calculated as

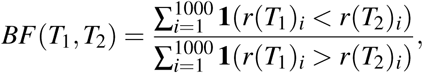

where *r*(*T*_1_)_*i*_ is the rank of team *T*_1_ on the *i*-th bootstrap, *r*(*T*_2_)_*i*_ is the rank of team *T*_2_ on the *i*-th bootstrap, and **1** is the indicator function.

### 4.2 Entropy and spatial autocorrelation

The entropy of a binarized *in situ* measurements of gene *G* was calculated as

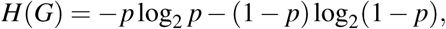

where p is the probability of gene G to have value 1. In other words, *p* is the fraction of cells where *G* is expressed.

The join count statistic is a measure of a spatial autocorrelation of a binary variable. We will refer to the binary expression 1 and 0 as black (*B*) and white (*W*). Let *n*_*B*_ be the number of bins where *G* is expressed (*G* = *B*), and *n*_*W*_ = *n - n*_*B*_ the number of bins where *G* is not expressed (*G* = *W*). Two neighboring spatial bins can form join of type *J* ∈ {*WW, BB, BW*}.

We are interested in the distribution of BW joins. If a gene has a lower number of BW joins that the expected number of BW, then the gene is positively spatially autocorrelated, i.e., the gene is highly clustered. Contrarily, higher number of BW joins points towards negative spatial correlation, i.e dispersion.

Following Cliff and Ord [27] and Sokal and Oden [28], the expected count of BW joins is

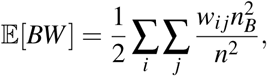

where the spatial connectivity matrix *w* is defined as

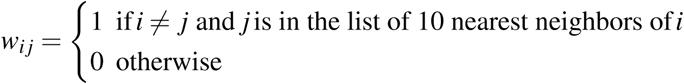

The variance of BW joins is

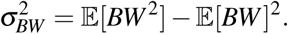

where the term 𝔼 [*BW* ^2^] is calculated as

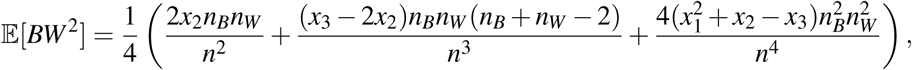

where 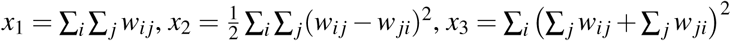.

Note that the connectivity matrix *w* can also be asymmetric, since it is defined by the nearestm neighbor function.

Finally, the observed BW counts are

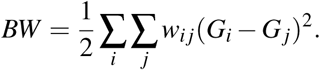

The join counts test statistic is then defined as

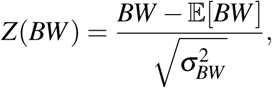

which is assumed to be asymptotically normally distributed under the null hypothesis of no spatial autocorrelation. Negative values of the *Z* statistic represent positive spatial autocorrelation, or clustering, of gene *G*. Positive values of the *Z* statistic represent negative spatial autocorrelation, or dispersion, of gene *G*.

### 4.3 Implementation details

The challenge scoring was implemented and run in R version 3.5, the post analysis was performed with R version 3.6 and the core `tidyverse` packages. We used the publicly available implementation of `DistMap` (https://github.com/rajewsky-lab/distmap). MCC calculated with R package `mccr` (0.4.4). t-SNE embedding and visualization produced with R package `Rtsne` (0.15). DBSCAN clutering with R package `dbscan` (1.1-4).

### 4.4 Code availability

https://github.com/dream-sctc/Scoring

### 4.5 Data description

#### Reference Database

The reference database comes from the Berkeley *Drosophila* Transcription Network Project. The *in situ* expression of 84 genes (columns) is quantified across the 3039 *Drosophila* embryonic locations (rows) for raw data and for binarized data. The 84 genes were binarized by manually choosing thresholds for each gene.

#### Spatial coordinates

One half of *Drosophila* embryo has 3039 cells places as x, y and z (columns) for a total of 3039 embryo locations (rows) and a total of 3039·3 coordinates.

#### Single cell RNA sequencing

The single-cell RNA sequencing data is provided as a matrix with 8924 genes as rows and 1297 cells as columns. In the raw version of the matrix, the entries are the raw unique gene counts (quantified by using unique molecular identifiers – UMI). The normalized version is obtained by dividing each entry by the total number of UMIs for that cell, adding a pseudocount and taking the logarithm of that. All entries are finally multiplied by a constant. For a given gene and only considering the Drop-seq cells expressing it we computed a quantile value above (below) which the gene would be designated ON (OFF). We sampled a series of quantile values and each time the gene correlation matrix based on this binarized version of normalized data versus the binarized BDTNP atlas was computed and compared by calculating the mean square root error between the elements of the lower triangular matrices. Eventually, the quantile value 0.23 was selected, as it was found to minimize the distance between the two correlation matrices.

The short sequences for each of the 1297 cells in the raw and normalized data are the cell barcodes.

## 5 Acknowledgments

This research was funded in part by PROACTIVE 2017 “From Single-Cell to Multi-Cells Information Systems Analysis” (C92F17003530005 Department of Information Engineering, University of Padova) for B.D.C.; National Institutes of Health grant number U54CA21729 for J.R.; ICMR JRF (Indian Council of Medical Research - Junior Research Fellowship) for S.A.and X.W. was funded by the National Natural Science Foundation of China (No.61702421 and No.61772426).

## 6 Author contributions

Conceptualization, N.K., N.R., J.S.R., G.S., and P.M.; Methodology, J.S., M.E.A., G.S., and P.M.; Software, J.T., and M.E.A.; Formal Analysis, J.T., M.E.A., G.S. and P.M.; Writing - Original Draft, J.T. and P.M.; Writing - Supervision, J.S.R., G.S., and P.M. - R.K, E.G. and P.B produced animated figures of results at https://dream-sctc.uni.lu/

## 7 Competing interests

The authors declare no competing interests.

## 8 Materials and Correspondence

Requests for data, resources, and or reagents should be directed to PabloMeyer.

## Notes

#### Summary of Updates

Supplementary materials were added

https://dream-sctc.uni.lu/

