## Supplementary materials for "Predicting cellular position in the *Drosophila* embryo from Single-Cell Transcriptomics data"

Giacomo Baruzzo[26], Marco Cappellato[26], Irene Zorzan[26], Simone Del Favero[26], Luca Schenato[26], Fabio Vandin[26], Barbara Di Camillo[26], Shruti Gupta[27], Ajay Kumar Verma[27], Shandar Ahmad[27], Ronesh Sharma[28,29], Edwin Vans[28,29], Alok Sharma[29,30,31,32], Ashwini Patil[33], Alejandra Carrea[34], Andres M. Alonso[35,36], Luis Diambra[35,36], Vijay Narsapuram[37], Vinay Kaikala[37], Chaitanyam Potnuru[37], Sunil Kumar[37], Jiajie Peng[38], Xiaoyu Wang[38], Xuequn Shang[38], Dani Livne[39], Tom Snir[39], Hagit Philip[39], Alona Zilberberg[39], Sol Efroni[39], Hamid Reza Hassanzadeh[40], Reihaneh Hassanzadeh[41], Ghazal Jahanshahi[42], M-Mahdi Naddaf-Sh[43], Phillip M. Drayer[43], Sadra Naddaf-Sh[44], MarouenBen Guebila[45], Changlin Wan[46], Yuchen Cao[47], Saber Meamardoost[48], Nan Papili Gao[49], and Rudiyanto Gunawan[48]

[26]Department of Information Engineering, University of Padova, Padova, Italy
[27]School of Computational & Integrative Sciences, Jawaharlal Nehru University, New Delhi, India
[28]School of Electrical and Electronics Engineering, Fiji National University, Suva, Fiji
[29]School of Engineering and Physics, University of the South Pacific, Suva, Fiji
[30]Laboratory for Medical Science Mathematics, RIKEN Center for Integrative Medical Sciences, Yokohama, Japan
[31]Institute for Integrated and Intelligent Systems, Griffith University, Brisbane, Australia
[32]CREST, JST, Tokyo, Japan
[33]Human Genome Center, The Institute of Medical Science, The University of Tokyo, Tokyo, Japan
[34]CREG, Universidad Nacional de La Plata, Argentina
[35]InTech, Universidad Nacional de San Martin, Argentina
[36]CONICET, Argentina
[37]Data Science and Informatics, Corteva Agrisciences, Ascendas IT Park, Madhapur, Hyderabad, India
[38]School of Computer Science, Northwestern Polytechnical University, Xi'an, China
[39]Bar-Ilan University, Ramat Gan, Israel
[40]School of Interactive Computing, Georgia Institute of Technology, Atlanta, GA, USA
[41]Department of Computer Science, Georgia State University, Atlanta, GA, USA
[42]J. Mack Robinson College of Business, Georgia State University, Atlanta, GA, USA
[43]Department of Electrical Engineering, Lamar University, Beaumont, TX, USA
[44]Computer Engineering Department, Ferdowsi University of Mashhad, Mashhad, Iran
[45]Department of Biostatistics, Harvard T.H. Chan School of Public Health, Boston, MA, USA
[46]Department of Electrical and Computer Engineering, Purdue University, West Lafayette, IN, USA
[47]Department of Statistics, Purdue University, West Lafayette, IN, USA
[48]Chemical and Biological Engineering Department, University at Buffalo, The State University of New York, Buffalo, NY, USA
[49]Institute for Chemical and Bioengineering, ETH Zurich, Zurich, Switzerland

### Results from the challenge

For all animated figures see: https://dream-sctc.uni.lu/

Subchallenge 1: Reconstruction of spatial location of cells using 60 genes.

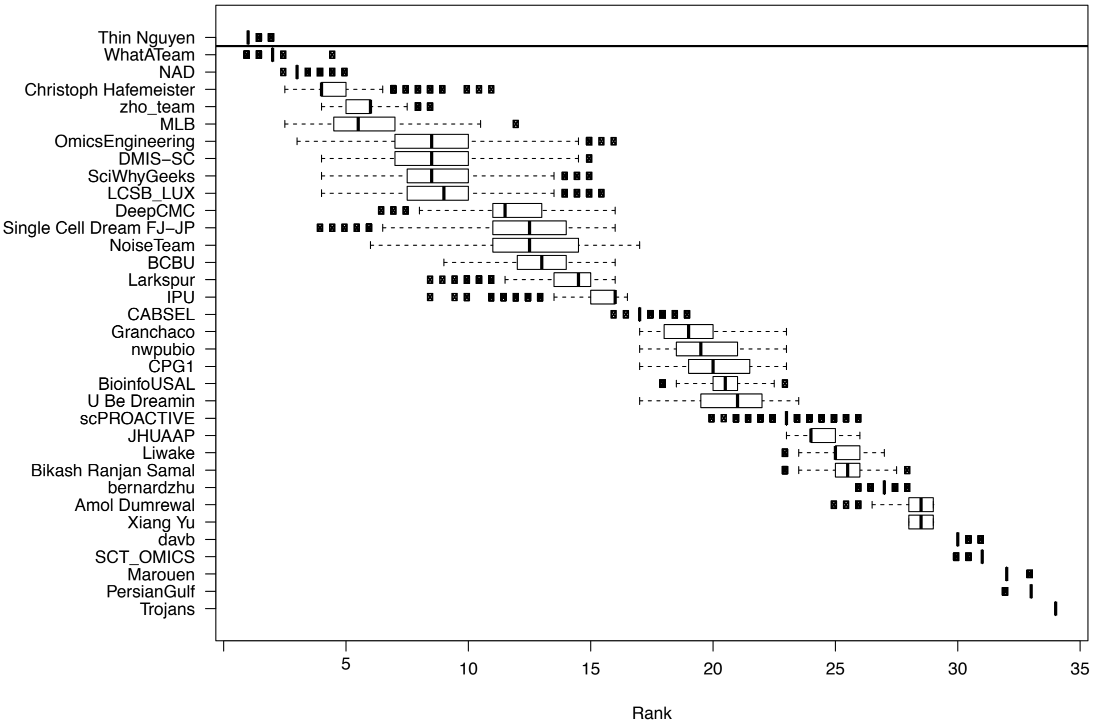


Figure S1: Results from the challenge showing boxplots of the average ranking across the 3 scoring schemes for the participating teams for 1000 bootstraps of the gold standard.

Subchallenge 2: Reconstruction of spatial location of cells using 40 genes.

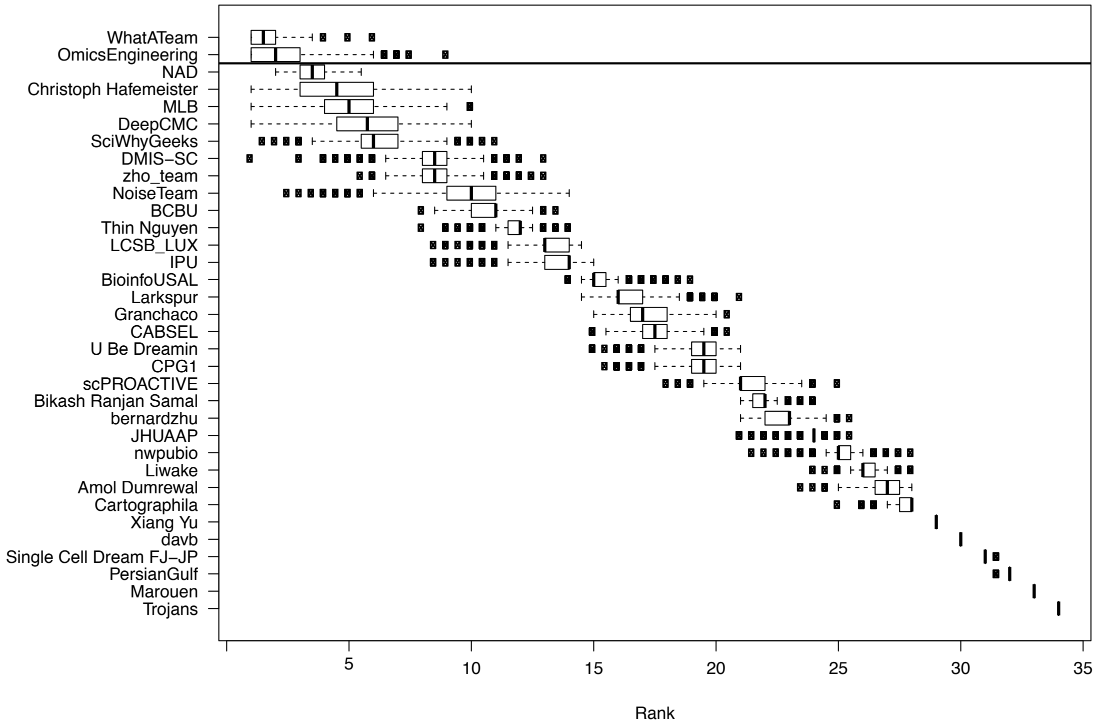


Figure S2: Results from the challenge showing boxplots of the average ranking across the 3 scoring schemes for the participating teams for 1000 bootstraps of the gold standard.

Subchallenge 3: Reconstruction of spatial location of cells using 20 genes.

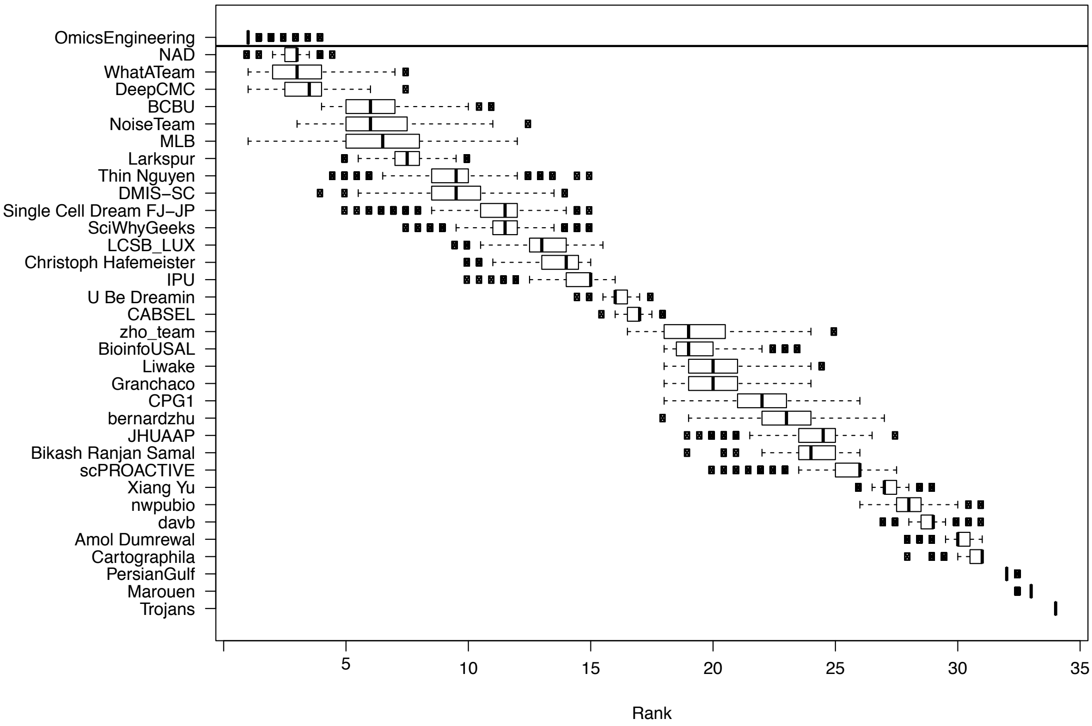


Figure S3: Results from the challenge showing boxplots of the average ranking across the 3 scoring schemes for the participating teams for 1000 bootstraps of the gold standard.

554 ## Additional figures and tables

Table S1: Methods used by the top 10 teams (ordered alphabetically) for gene selection and location prediction. The methods used for gene selection are categorized in four different categories: SFR - Supervised feature ranking, UFR - unsupervised feature ranking, KNW - background knowledge, and VAR - variance. The methods used for location3 prediction are categorized in three different categories: CMB - Combination of model prediction and MCC, MCC - Matthews correlation coefficient, and SIM - Similarity measure (non MCC). More detailed description (writeup) of the methods used by each team is available in the Leaderboards section on the challenge webpage `https://www.synapse.org/#!Synapse:syn15665609/wiki/583233`.

| Team | Selection | Prediction |
|---|---|---|
| BCBU | SFR - Random Forest | CMB - Random Forest, MCC |
| Challengers18 | UFR - Particle Swarm Optimization | SIM - Weighted correlation |
| Christoph Hafemeister | UFR - PCA (principal component analysis) on most variable genes, Expression correlation | MCC |
| Christoph Hafemeister | UFR - PCA on most variable genes, Expression correlation | SIM - Correlation |
| DeepCMC | SFR - LASSO (least absolute shrinkage and selection operator) | MCC |
| DeepCMC | SFR - Neural Network | MCC |
| MLB | UFR - Stepwise regression, PCA, k-nearest neighbors, F-score | SIM - F-score |
| NAD | SFR - Feedforward neural network, KNW - Clustering, VAR | CMB - Feedforward neural network, MCC |
| OmicsEngineering | UFR - Euclidean distance of expression | MCC |
| OmicsEngineering | SFR - Random Forest, Genetic algorithm | MCC |
| Thin Nguyen | VAR | MCC |
| Thin Nguyen | UFR - Nonnegative Discriminative Feature Selection | SIM - k-nearest neighbors |
| WhatATeam | KNW, Clustering | CMB - Local outlier factor, MCC |
| WhatATeam | UFR - Stepwise regression | CMB - Local outlier factor, MCC |
| Zho | UFR - Hierarchical clustering | SIM - Hamming distance, Silhouette score |

Table S2: Summary of methods used by the top 10 teams for gene selection and location prediction. Some teams used different approaches or a combination of approaches for different subchallenges. The categories of the method used for gene selection and location prediction are the same as in Table S1.

| | | Selection | | | |
|---|---|---|---|---|---|
| | | SFR | UFR | KNW | VAR |
| Prediction | CMB | 2 | 1 | 2 | |
| | MCC | 3 | 2 | | 1 |
| | SIM | | 5 | | |

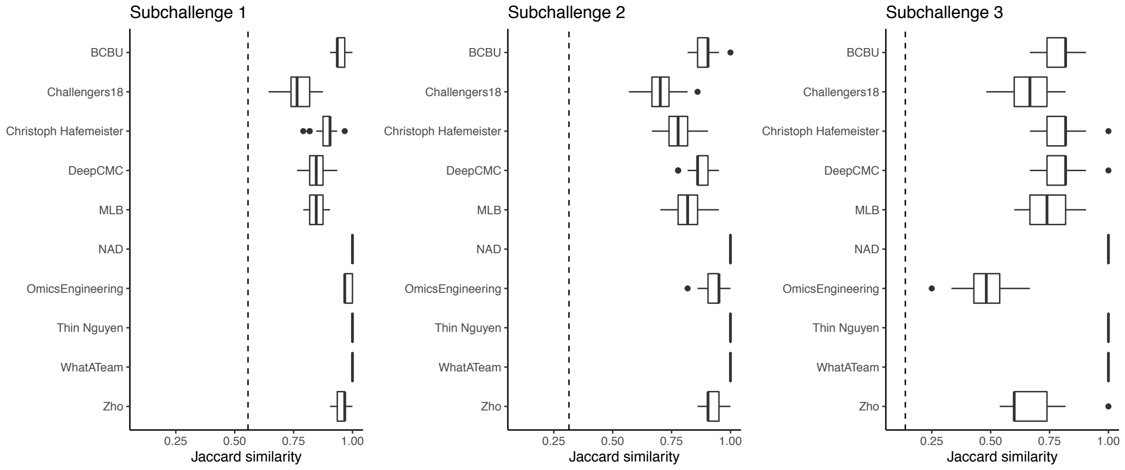


Figure S4: Boxplots of the Jaccard similarity between the genes selected for each of the 10 CV scheme in all 3 subchallenges. The teams that used the statistical properties of the genes as selection criteria, for example maximum variance, selected the same set of genes for all folds. This is expected since the distribution of a random subsample was selected to have the same properties as the original sample. Dotted line represents the limit for significance, i.e., the expected Jaccard similarity between two sets of randomly selected 60, 40 or 20 genes.

Table S3: Most frequently selected 60, 40 and 20 genes in subchallenges 1,2 and 3 respectively, in alphabetical order, colored according to Figure 5 from the main text. That is yellow are gap genes and green are pair-rule genes

| | |
|---|---|
| Subchallenge 1 | *aay ama ance antp apt blimp-1 brk btk29A bun cg10479 cg14427 cg43394 cg8147 croc cyp310a1 d dan danr dfd disco doc2 doc3 dpn E(spl)m5-HLH edl eve fj fkh ftz gt h hb htl ilp4 impE2 impL2 kni knrl kr lok mdr49 mes2 mESR3 noc nub oc odd prd rau rho run sna srp tkv toc traf4 trn tsh twi zen zfh1* |
| Subchallenge 2 | *aay ama ance antp blimp-1 brk btk29A cg43394 cg8147 croc cyp310a1 d dan disco doc3 dpn edl fj fkh ftz gt h ilp4 impE2 impL2 kni knrl kr mes2 mESR3 noc nub oc rho run sna srp tsh twi zfh1* |
| Subchallenge 3 | *ama antp brk cg8147 cyp310a1 disco doc2 doc3 fkh h ilp4 impE2 kni knrl mes2 nub oc sna tsh twi* |

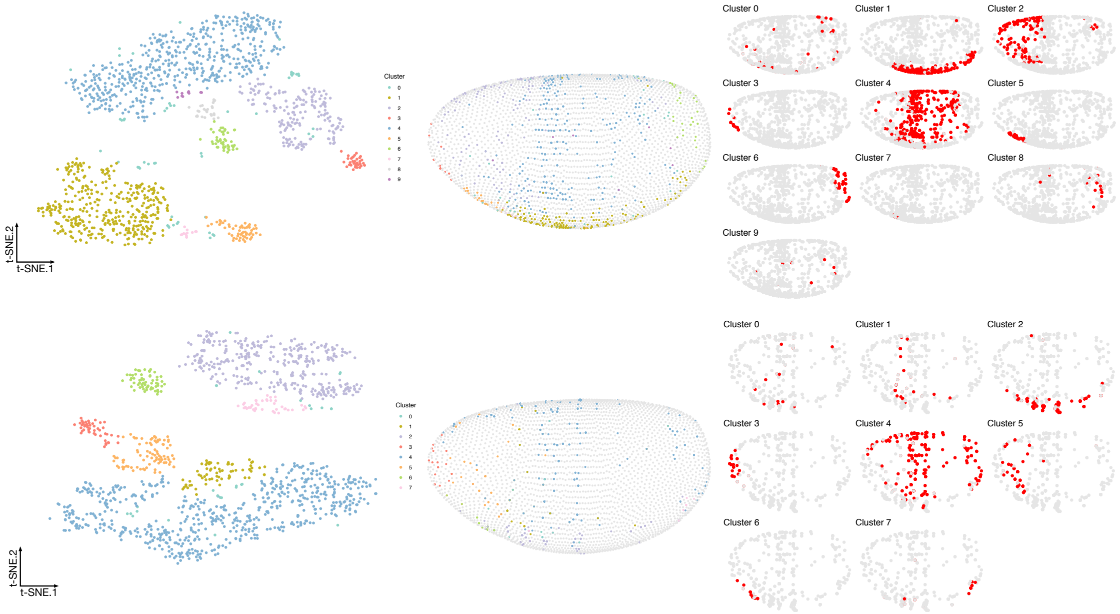


Figure S5: Visualization of the transcriptomics data containing only the most frequently selected **A** 40 genes from subchallenge 2 and **B** 20 genes from subchallenge 3 by the top performing teams (embedding to 2D by t-SNE). *Left* each point (cell) is filled with the color of the cluster that it belongs to (density-based clustering with DBSCAN). *Middle*, spatial mapping of the cells in the Drosophila embryo as assigned by DistMap using only the 60 most frequently selected genes from subchallenge 1. The color of each point corresponds to the color of the cluster from the t-SNE visualization. *Right*, highlighted (red) location mapping of cells in the Drosophila embryo for each cluster separately.
